# Multisensory integration in neurons of the medial pulvinar of macaque monkey

**DOI:** 10.1101/2022.03.21.485176

**Authors:** Anne-Laure Vittek, Cécile Juan, Lionel G. Nowak, Pascal Girard, Céline Cappe

**Affiliations:** Centre de Recherche Cerveau et Cognition (CerCo), CNRS UMR 5549, Université de Toulouse, UPS, Toulouse, France; INSERM, Toulouse, France

**Keywords:** multisensory, medial pulvinar, audiovisual, visual, auditory, single-unit

## Abstract

The pulvinar is a heterogeneous thalamic nucleus, which is well developed in primates. One of its subdivisions, the medial pulvinar, is connected to many cortical areas, including the visual, auditory and somatosensory cortices, as well as with multisensory areas and premotor areas. However, except for the visual modality, little is known about its sensory functions. A hypothesis is that, as a region of convergence of information from different sensory modalities, the medial pulvinar plays a role in multisensory integration. To test this hypothesis, two macaque monkeys were trained to a fixation task and the responses of single-units to visual, auditory and auditory-visual stimuli were examined. Analysis revealed auditory, visual and multisensory neurons in the medial pulvinar. It also revealed multisensory integration in this structure, mainly subadditive and suppressive. These findings show that the medial pulvinar is involved in multisensory integration.

## Introduction

Our perception of the environment is multisensory. Stein and Meredith were among the firsts to find multisensory neurons while recording in the superior colliculus (Meredith and Stein, 1986). Single-units recorded in this subcortical structure enabled them to define the rules of multisensory integration. Associative cortical areas of the temporal, frontal and parietal lobes were later shown to have multisensory properties. For example, the superior temporal sulcus, the orbitofrontal cortex and area 7 receive inputs from many unisensory cortical areas and individual neurons in these regions can respond to stimuli from several sensory modalities (Barraclough et al., 2005; Benevento et al., 1977; Bruce et al., 1981; Falchier et al., 2011 for a review; Leinonen et al., 1979). More recently, evidence that even unisensory cortical areas have multisensory properties has been reported (Cappe et al., 2009a for a review): anatomical connections between unisensory areas of different modalities have been observed (Cappe and Barone, 2005; Falchier et al., 2002, 2011). Moreover, studies showed that the responses of neurons in the auditory (Kayser et al., 2008; Lakatos et al., 2007), visual (Wang et al., 2008) and somatosensory (Zhou and Fuster, 2004) cortices can be modulated by the addition of another modality. All these results led Ghazanfar and Schroeder to the provocative proposition that the whole cortex might be multisensory (Ghazanfar and Schroeder, 2006).

However, besides cortex, subcortical structures may also be involved in multisensory integration. Among these, the medial pulvinar appears as an ideal candidate. Located posteriorly in the thalamus, the medial pulvinar presents reciprocal connections with unisensory cortical areas (auditory, visual and somatosensory cortices) (Cappe et al., 2007, 2009a), polysensory areas (superior temporal polysensory cortex, prefrontal cortex, posterior cingulate cortex, insular cortex) (Gutierez et al., 2000; Romanski et al., 1997) and motor areas (dorsal premotor cortex) (Morel et al., 2005). It also receives afferents from sub-cortical structures such as the superior colliculus (Trojanowski and Jacobson, 1975) and is interconnected with the amygdala (Jones and Burton, 1976). These numerous connections suggest that it is a region of convergence of information from different sensory modalities. The functions of the pulvinar are still under investigations (Froesel et al., 2021 for a review). First, lesion studies in humans (Snow et al., 2009; Ward and Arend, 2007; Wilke et al., 2018) and inactivation studies in monkeys (Wilke et al., 2010) showed deficits in visual attention and distractor filtering as well as deficits in localizing visual targets and in the control of visually-guided eye and limb movements (grasping and reaching). However, these behavioral studies did not examine sensory modalities other than visual. Second, the pulvinar is involved in the communication between cortical areas (Benarroch, 2015; Saalmann and Kastner, 2011). Third, the pulvinar has a role in sensory processing: electrophysiological studies showed that neurons in the pulvinar respond to auditory (Gattass et al., 1978; Mathers and Rapisardi, 1973; Yirmiya and Hocherman, 1987), somatosensory (Mathers and Rapisardi, 1973), and visual stimuli (Almeida et al., 2015; Benevento and Miller, 1981; Gattass et al., 1978; Komura et al., 2013; Maior et al., 2010; Mathers and Rapisardi, 1973; Nguyen et al., 2013; Van Le et al., 2013, 2014, 2016; Wilke et al., 2009). However, the subdivisions of the pulvinar in which the recordings were performed were not precisely identified in these studies. Furthermore, only one study (Gattass et al., 1978) tested stimuli of different modalities on the same neurons, yet did not examine multisensory interaction.

In the present study, we recorded single-units in a well-defined subdivision of the pulvinar, the medial pulvinar, of two awake macaques performing a fixation task. Single-unit data confirmed the presence of auditory, visual and audiovisual neurons in the medial pulvinar. Furthermore, it revealed the existence of a new type of multisensory neurons, which we named “complex multisensory”. We also provide evidence of multisensory integration in this structure, which is mostly sub-additive and suppressive.

## Results

### 1. Medial pulvinar contains auditory, visual and audiovisual neurons

We recorded spiking activity in response to visual, auditory and audiovisual stimuli in the medial pulvinar of two macaques engaged in a fixation task (figure 1). 374 single-units were recorded in 164 sites and 328 analyzed for responsiveness to the stimuli (46 neurons were excluded because of too low activity or fluctuating baseline). Most (213/328, 65%) medial pulvinar neurons responded to auditory, visual and/or audiovisual stimuli. These single-units were all spontaneously active, with a median spontaneous firing rate of 3.9 spikes/s. During stimulus presentation, the neurons either showed no change in firing rate (no response), an increase of firing rate (mean + 4.2 spikes/s relative to spontaneous activity level, global population response fig 2.A) or a decrease in activity (mean – 1.2 spikes/s, global population response fig 2.B).

**Figure 1:**
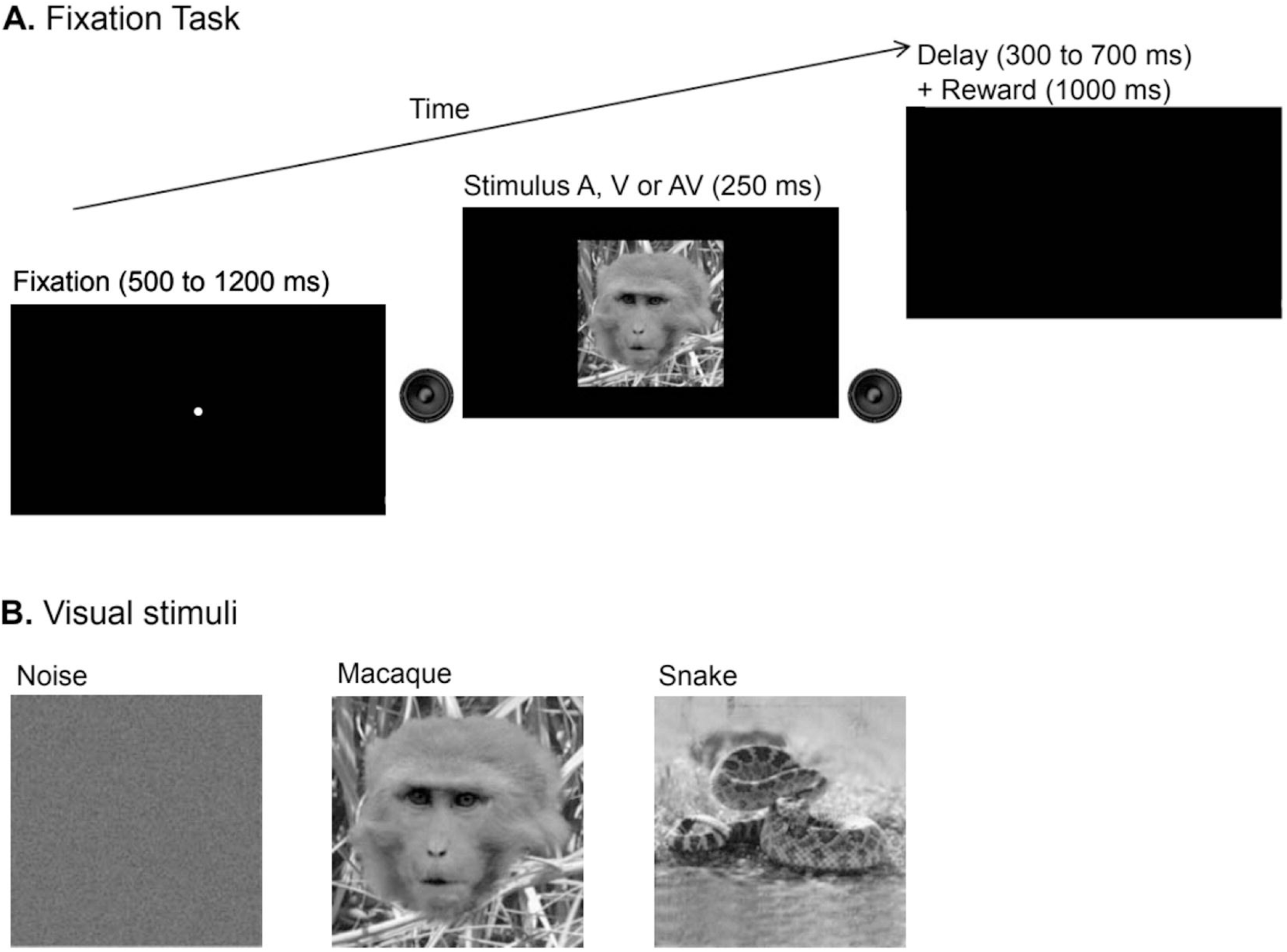
experimental design. **A) Schematic representation of the behavioral task.** The monkey sat in a primate chair in front of a computer screen. After fixation of the central point (during a random time between 500 and 1200 ms), an auditory, visual or audiovisual stimulus was presented for 250 ms. The monkey had to maintain fixation during stimulus presentation to obtain a reward (compote drop). **B) Visual stimuli**. Three static visual stimuli were presented (always the same picture for each): a picture of random dots, a macaque face and a rattlesnake.

**Figure 2:**
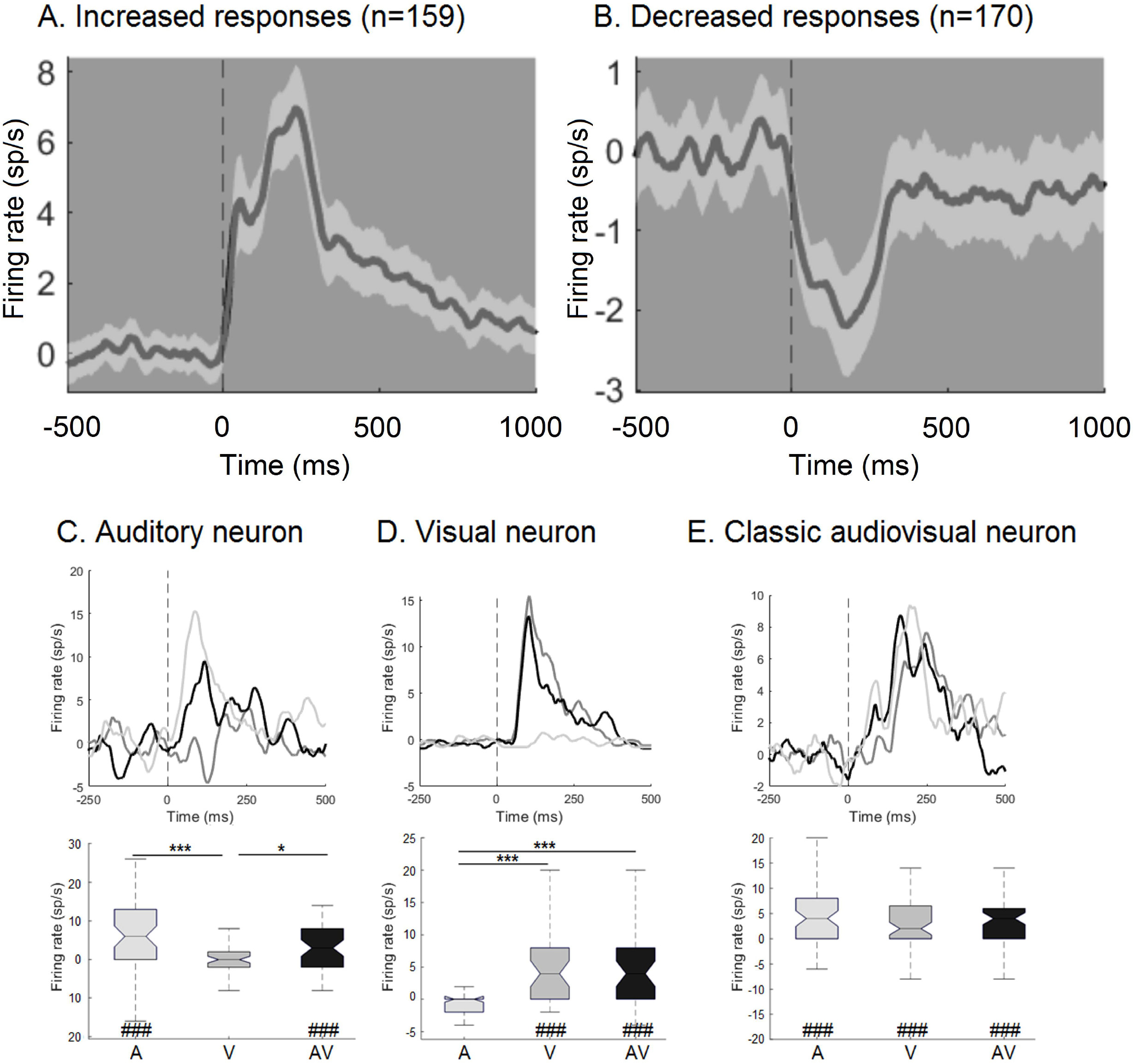
Global responses and individual neuron examples. A-B) Global PSTHs representing the mean firing rate (sp/s) of neuronal populations according to the response types: enhanced (A) or decreased (B) responses. Responses were averaged over all stimuli and all modalities. Black line corresponds to the mean and gray shading to ± 95% CI. C-E) Examples of an auditory neuron (C), a visual neuron (D) and a classic audiovisual neuron (E). Top: peri-stimulus time histograms (PSTHs) were calculated across all trials (all stimulus categories pooled) for a given modality (auditory in light gray, visual in gray and audiovisual in black), with a bin width of 10 ms and a 45 ms smoothing. Bottom: boxplots summarizing distributions of mean firing rates for each stimulus presentation in response to auditory (A), visual (V) and audiovisual (AV) stimuli. Paired spontaneous activity measured during 500 ms before stimulus onset was subtracted from each response. Responses were considered as significant when the firing rates during stimulus presentation were different from the preceding spontaneous firing rates, as tested with a Wilcoxon signed rank test. Sharps below boxplots indicate significance level: #: p<0.05, ##: p<0.01 and ###: p<0.001. Modulation of response amplitude by the modalities was evaluated by the Kruskal-Wallis test, followed by Mann-Whitney tests between modality (p-values adjusted by Bonferroni correction). Asterisks above boxplots indicate significance level: *: p<0.05, **: p<0.01 and ***: p<0.001.

The 213 responsive neurons were individually found to be auditory, visual, audiovisual or nonspecific to one modality. Unisensory cells responded to only one sensory modality, and responded identically to audiovisual stimuli. The two neurons presented in figure 2C, D are unisensory cells (2.C auditory, 2.D visual). For the neuron presented in figure 2.C, the auditory stimuli induced an increase in activity that was statistically significant when compared to baseline activity. Similarly, there was a statistically significant increase in activity in response to auditory-visual stimuli, but the activity level was not statistically different from the one obtained in response to auditory stimuli alone. Presentation of visual stimuli had no significant effect on activity. The neuron presented in figure 2.D has similar characteristics, but for visual stimuli: the activity increased by the same amount in response to visual and auditory-visual stimuli and did not change in the presence of auditory stimuli. At the population level, the medial pulvinar appears to have slightly more auditory than visual neurons (χ^2^=4,45, df=1, p=0.03): among the 213 responsive neurons, 59 (28%) were classified as auditory, and 39 (18%) as visual (figure 3.A).

**Figure 3:**
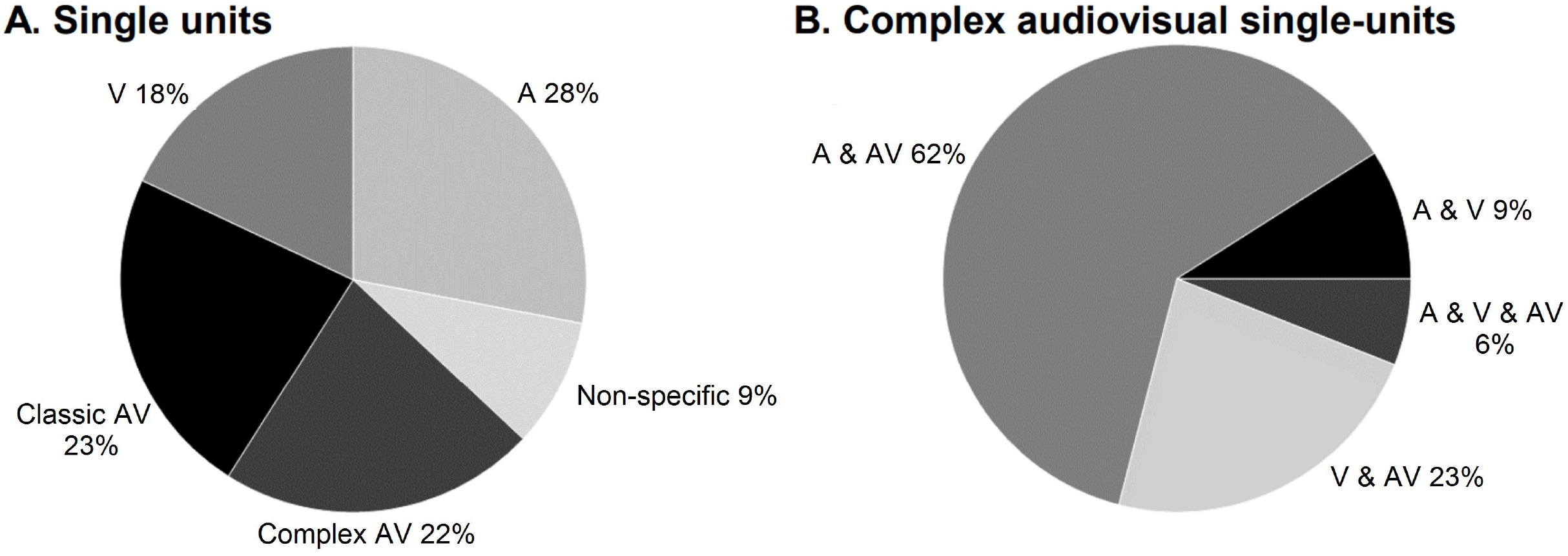
proportions of single-units classified by sensory modalities and multimodal integration types. A) Pie chart representing the proportions of single-units classified as visual only, auditory only, classic audiovisual, complex audiovisual or non-specific to a modality. B) Pie chart representing the proportions of complex audiovisual neurons, classified according to the modalities they responded to.

We found 49 “classic” audiovisual neurons (23% of the responsive neurons, fig 3A). These neurons could be classified in four classes, depending on their response features. The first class includes neurons that responded only to auditory-visual stimuli. This represents 82% of the classic audiovisual neurons in our sample. The second class, which represents 10% of the classic audiovisual neurons, groups cells which responded in all sensory conditions; an example is presented in figure 2.E. For this neuron, the visual, auditory and audiovisual stimuli induced an increase in activity that was statistically significant when compared to baseline activity. The third class corresponds to neurons that responded to auditory and visual stimuli but not to bimodal stimuli, and represents 6% of the classic audiovisual neurons. Finally, some neurons responded to a single modality when tested with unimodal stimuli, but the response was different from that to the audiovisual stimuli. This fourth category represents 2% of our sample.

### 2. Medial pulvinar contains complex audiovisual neurons

47 neurons were found to possess “complex” audiovisual responses (22% of the responsive neurons, fig 3A). These are cells for which the sensory modality that evoked a response depended on the stimulus category. For example, the neuron in figure 4 presented a statistically significant increase in activity in response to the noise stimulus in both visual, auditory and audiovisual modalities, to the snake stimulus also in visual, auditory and audiovisual modalities, but to the macaque stimulus only in the visual and audiovisual modalities. This cell did not respond to the auditory macaque stimulus but responded equally well to the visual and audiovisual macaque stimuli. It was classified as visual audiovisual complex multisensory neuron, like 23% of the complex audiovisual neurons. We found three other types of complex audiovisual neurons (figure 3B): auditory audiovisual (62% of the neurons), auditory visual (9%) and auditory visual audiovisual (6%) (figure 3.B).

**Figure 4:**
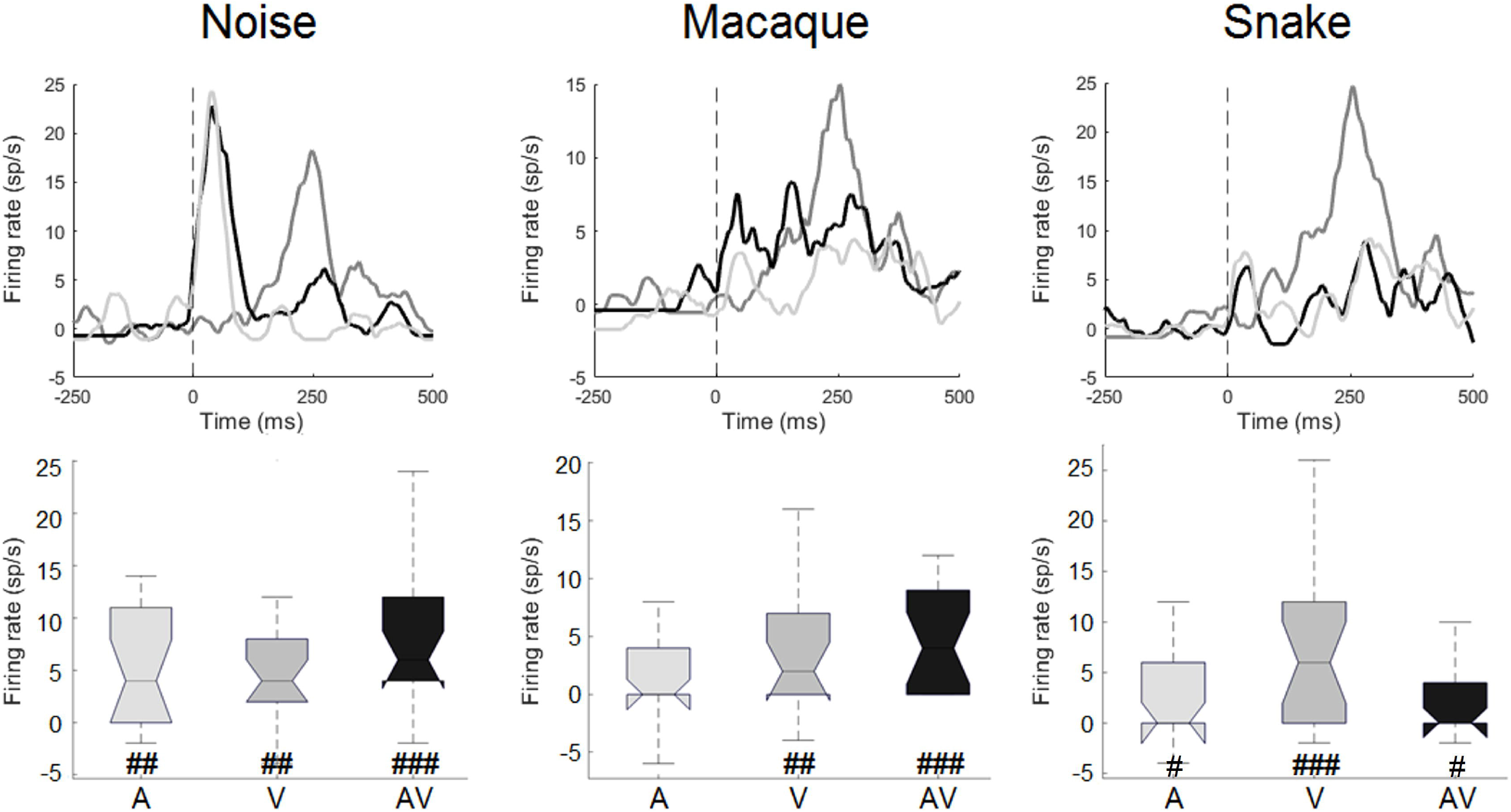
example of a complex audiovisual neuron. First column: responses to noise stimuli; second column: responses to macaque stimuli; third column: responses to snake stimuli. Boxplots summarizing the firing rate distributions for each stimulus are shown below each corresponding PSTH (auditory in light gray, visual in gray and audiovisual in black). This complex neuron was assigned to the visual modality for macaque stimulation and to the multisensory modality for noise and snake stimuli. Significance of responses were tested using Wilcoxon test (#: p<0.05, ##: p<0.01 and ###: p<0.001) and modality modulation by Mann Whitney test (p-values adjusted with the Bonferroni correction; *: p<0.05, **: p<0.01 and ***: p<0.001).

### 3. Medial pulvinar neurons are poorly selective

The sparseness and the selectivity indexes indicate that medial pulvinar neurons responded to many stimuli and tended to be poorly selective. The sparseness index was computed using the formula:

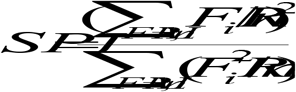

with FR_i_ the response to the stimulus i, and n=9 (number of stimuli). It quantifies the variability of the responses to all nine stimuli used. It is indicative of a sparse representation when the index approaches 0 and of a distributed representation when the index reaches values close to 1. In our sample, it ranges from 0.42 to 1, with a median at 0.93 (figure 5).

**Figure 5:**
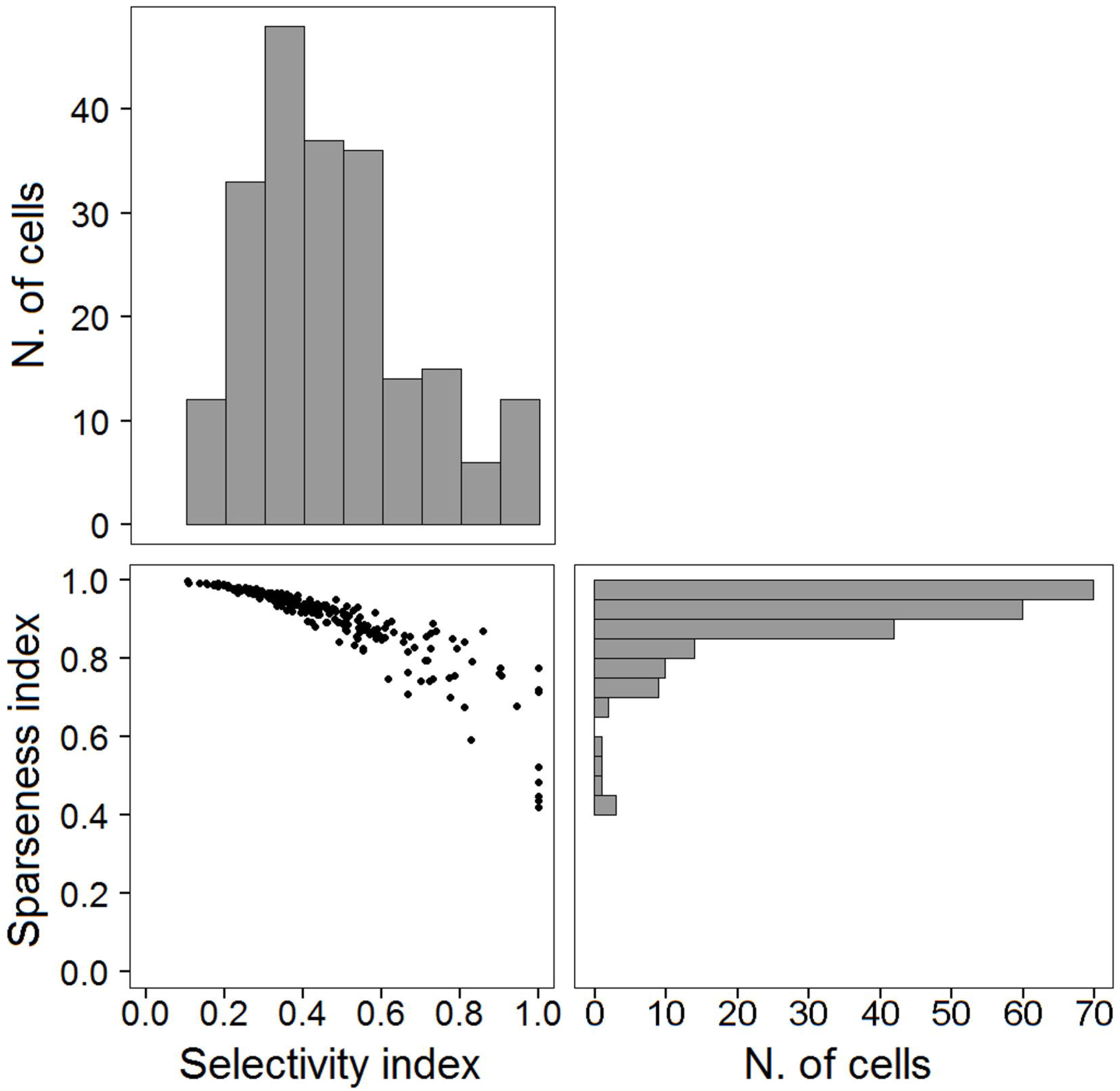
neuronal selectivity and sparseness. The sparseness indexes of the 213 responding cells are plotted as a function of the selectivity indexes (bottom left). Indexes distributions are shown on the top (selectivity index) and on the right (sparseness index).

The selectivity index compares the best and the worst responses of a neuron. Its formula is:

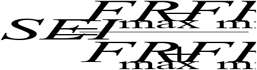

with FR_min_ and FR_max_ the minimal and maximal responses among the nine responses. An index close to 1 is indicative of a very selective neuron, which responded to only one stimulus, whereas an index close to 0 is indicative of a poorly selective neuron, which responded almost equally well to all stimuli. The selectivity index ranges from 0.10 to 1, with a median at 0.43 (figure 5).

In addition, population responses were not different between snake, macaque and noise stimuli, whatever the modality ((all p-values > 0.05) (figure S1 for PSTH according to stimuli, modalities and response types).

### 4. Multisensory integration in the medial pulvinar is sub-additive and suppressive

Additivity and amplification indexes, calculated for each multisensory cell, revealed sub-additive and suppressive multisensory integration in the medial pulvinar. The additivity index compares the audiovisual response to the sum of the unisensory responses. The formula of this index is:

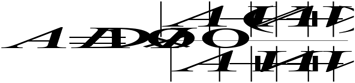

with AV, V and A the median of the mean firing rates in the audiovisual, visual and auditory conditions. It informs about sub-additive (index<0), additive (index=0) or supra-additive (index>0) interactions between these responses. The amplification index, comparing the audiovisual response to the best unisensory response, allows determining if the multisensory response is enhanced (index>0) or suppressed (index<0) compared to the best unisensory response. The formula of this index is:

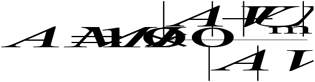

with U_max_ the median of the mean firing rate in the best unisensory condition.

Additivity index ranges between −0.74 and 0.13 (94 of the 96 multisensory neurons have a negative index) and amplification index ranges between −0.61 and 0.34 (61 of the 96 multisensory neurons have a negative index). There is a continuum from sub-additive and depressed to supra-additive and enhanced responses (figure 6) and the majority of the cells are sub-additive depressed cells.

**Figure 6:**
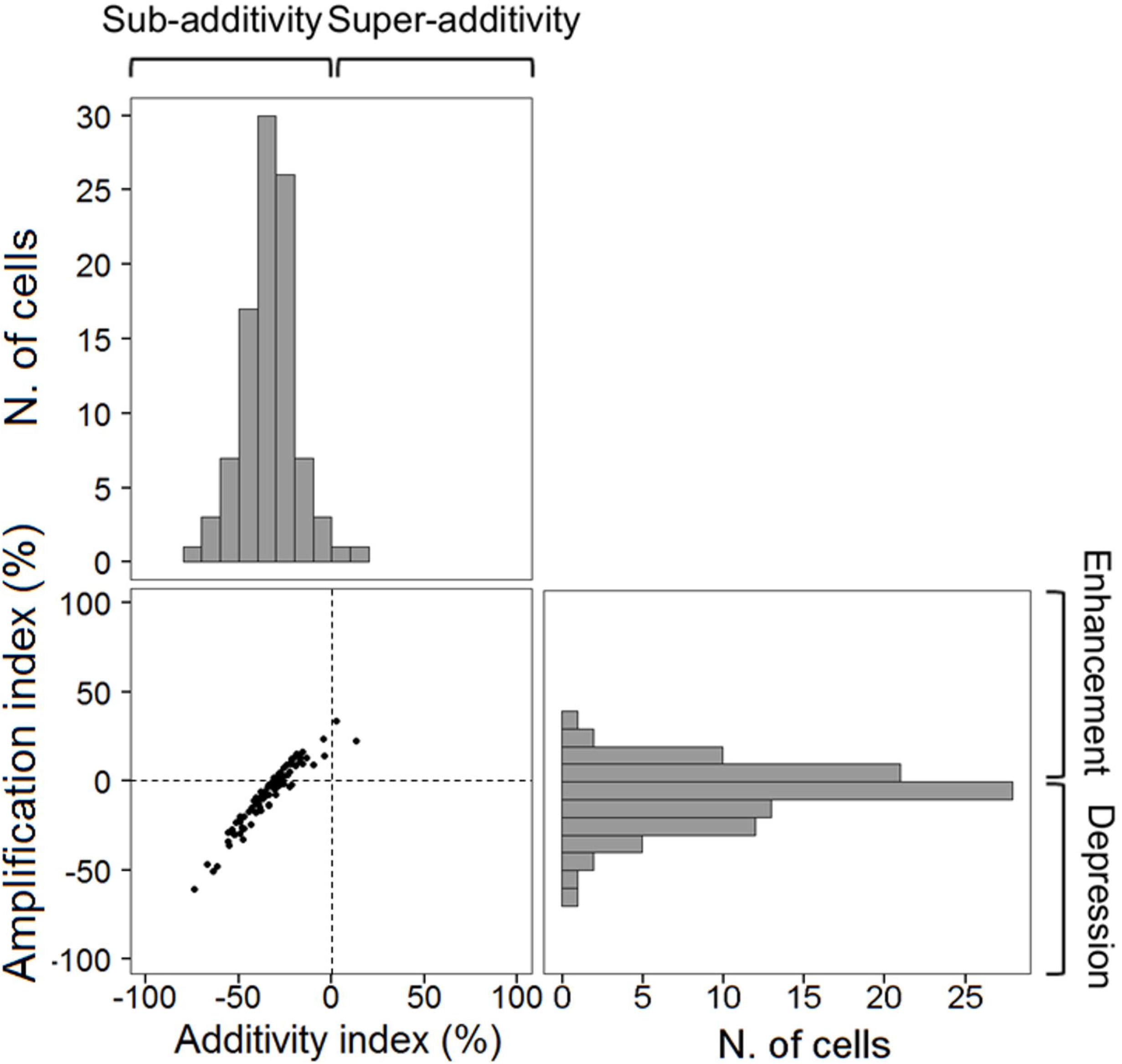
multisensory integration: additivity and amplification indexes. The amplification indexes of 96 multisensory cells are plotted as a function of additivity indexes (bottom left). Indexes distributions are shown on the top (additivity index) and on the right (amplification index).

### 5. Faster responses for auditory and audiovisual than for visual stimuli

The latencies of the responses were measured as the half rise latency (see methods). Latencies have been obtained for 69 neurons, yielding a total of 160 latencies as a given neuron could respond to several stimulus categories and modalities. We first examined the influence of stimulus modality. For this analysis we reduced the dataset by taking the shortest latency within each modality whenever latencies were available for more than one stimulus category. Stimulus modality strongly affected latencies (Kruskal-Wallis test, P<0.0001). The medians of the half-rise latencies for auditory and audiovisual responses were 89 ms (n=35) and 75 ms (n=46), respectively. The distributions for auditory and audiovisual latencies largely overlapped and did not differ significantly (Posthoc Dunn’s test, P>0.99). On the other hand, visual latencies were longer than both auditory (P=0.003) and audiovisual latencies (P<0.0001). The cumulative distribution shows a systematic lag for visual responses by about 50 ms, with a median visual latency at 141 ms (n=30) (figure 7).

**Figure 7:**
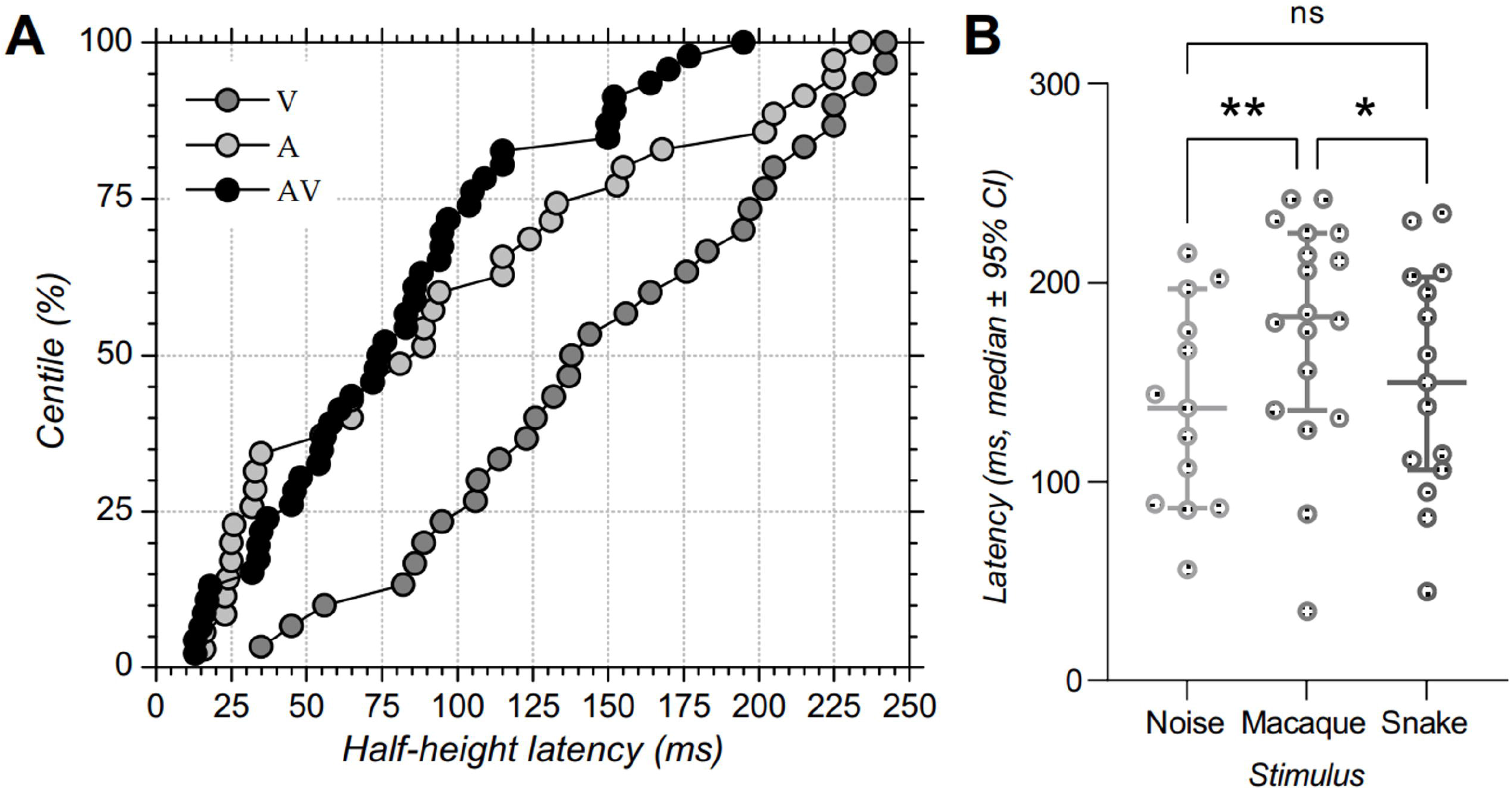
latencies of medial pulvinar neurons. A. Cumulative distribution of half-rise latencies for each modality: visual (V, 30 latencies) in gray, auditory (A, 35 latencies) in light gray and audiovisual (AV, 46 latencies) in black. B. Distribution of half-rise latencies for visual responses produced by the presentation of noise, macaque and snake pictures. Dots represent individual data, horizontal central bars corresponds to the medians and error bars indicate the 95% confidence interval of the medians. Latencies were significantly longer for responses to macaque stimuli. Asterisks indicate significance level obtained with Posthoc Tukey’s test: *: p<0.05; **: p<0.01.

Latencies were also compared between stimuli in each modality. These comparisons did not reveal differences for the auditory and audiovisual modalities (mixed effect model, P=0.13 and 0.09 respectively). However, as summarized in figure 7B, differences between stimuli were observed in the visual modality (P=0.01) and revealed longer latencies for macaque stimulus (median 183 ms, n=18) compared to noise (median 137 ms, n=13) and snake (median 150 ms, n=15) (Posthoc Tukey’s test: P=0.002 for comparison between macaque and noise, P=0.02 between snake and macaque, P=0.06 between noise and snake).

### 6. Sensory modalities are intermingled in the medial pulvinar

A 3-dimensional map was constructed to report the location of the recording sites in the explored region (2D representation in figure 8). No topographic organization is visible for the sample of recorded neurons: neurons of all categories were present in variable proportions at each recording site. This was confirmed by a PCA in 3D space by using the three dimensions of recording site coordinates (AP, ML and depth).

**Figure 8:**
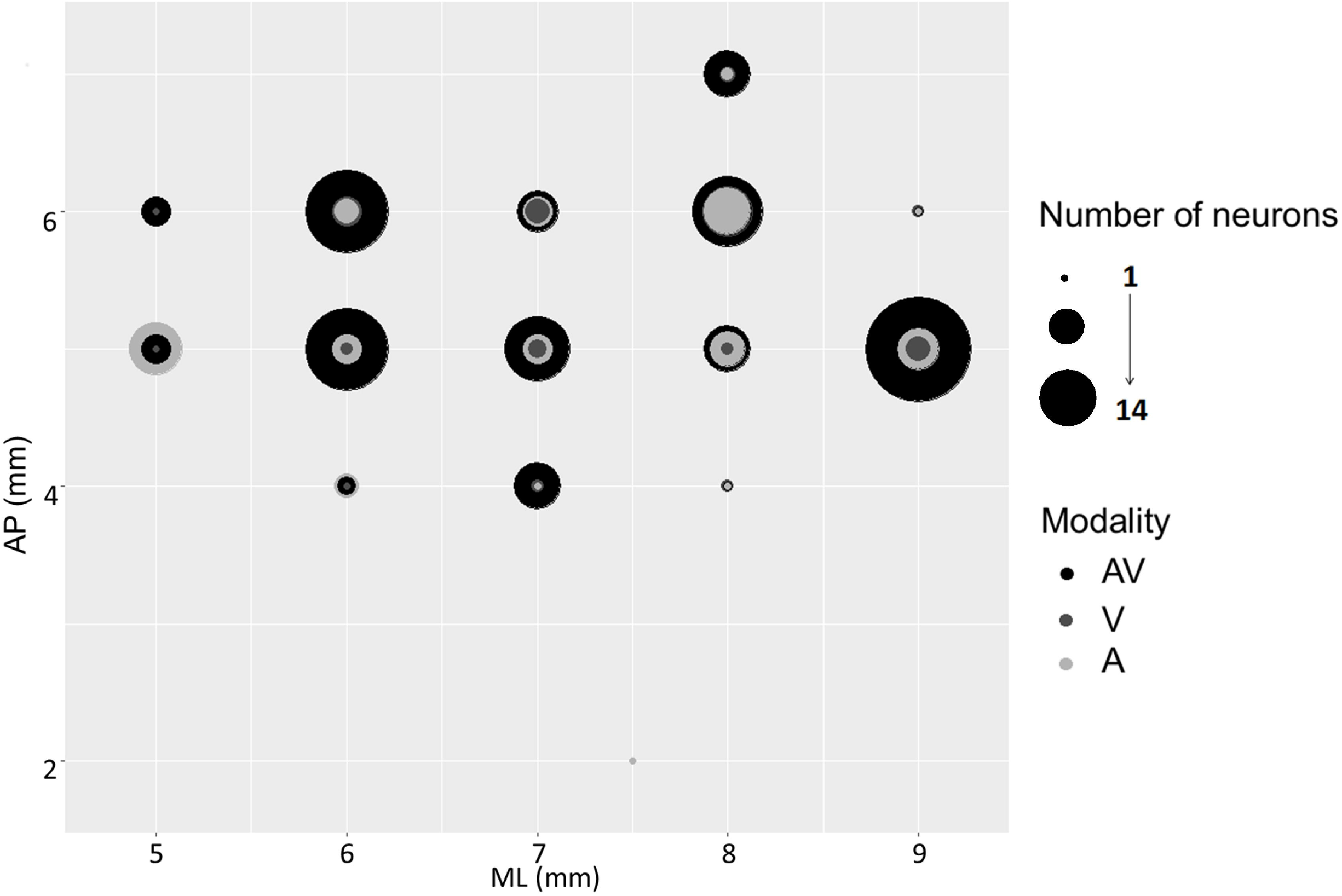
sensory modalities are intermingled in the medial pulvinar. Topographic representation of neuronal responses to each modality. Representation of response modalities of the 213 cells with significant responses as a function of the anteroposterior (y axis, from 2 to 6 mm) and mediolateral (x axis, from 5 to 9 mm) coordinates of electrode penetration sites. Response modality is gray-scale-coded (A in light gray, V in dark gray and AV in black). The radius of the circles is proportional to the number of neurons that responded to each modality.

## Discussion

### Unisensory and multisensory neurons in the medial pulvinar

In the present study, we investigated neuronal responses to auditory, visual and audiovisual stimuli in the medial pulvinar. Even though numerous studies have tested visual responses in the pulvinar (Mathers and Rapisardi, 1973; Van Le et al., 2016), only a few have focused on the medial pulvinar, or used auditory stimuli (i.e. Yirmiya and Hocherman, 1987). We confirmed that neurons in the pulvinar are responsive to visual stimulations and auditory stimulations. We also reported multisensory audiovisual neurons and we described and quantified multisensory integration in the medial pulvinar of awake monkeys.

Another novelty in our results lies in the identification of complex audiovisual neurons. The modality to which these neurons responded depended on the stimulus category. This type of neurons has not been described previously. Indeed, identifying these neurons requires different types of multisensory stimuli. Instead, previous studies most often relied on the use of a single multisensory stimulus. The existence of complex audiovisual neurons is consistent with multidimensional coding, as suggested by Chang and Tsao for facial identity (Chang and Tsao, 2017). In this hypothesis, responses of neurons can be positioned in a space with many dimensions relevant to the used stimuli. Thus, neurons are not responsive to the stimuli but to some of their constitutive features. Features can be common to several stimuli (for example, visual noise and visual snake may share common features), but are not necessarily shared by the auditory and visual versions of a given stimulus. Thus, a neuron can respond to some visual stimuli and to different auditory stimuli, depending on which stimuli share the features to which it responds. This corresponds to our complex audiovisual stimuli. Using more stimuli would help to observe more complex neurons, and understand better what are the relevant features, and how the modalities play a part in this multidimensional coding. The pulvinar could be viewed as a detector of incoming stimuli without *a priori* about any category. If one supposes population coding of these stimuli, this could lay on a multidimensional description of these stimuli with auditory and visual components belonging to different categories.

### Multisensory integration in the medial pulvinar

The presence of numerous audiovisual neurons prompted us to examine their multisensory integration properties. Multisensory integration was globally sub-additive (additivity index < 0) and suppressive (amplification index < 0). The first studies on multisensory integration, performed in the superior colliculus of the cat and monkey, found mainly enhanced responses (Meredith and Stein, 1986; Perrault Jr et al., 2003). More recent studies, based on recordings in the superior colliculus (Populin and Yin, 2002), the ventral intraparietal area (Avillac et al., 2007), the auditory cortex (Kayser et al., 2008) and the ventrolateral prefrontal cortex (Sugihara et al., 2006), mainly found suppressive and sub-additive responses. In addition to the recording sites or the species, the task and the state (anesthetized vs. awake) of the animals should be considered when comparing results. In the studies of Meredith and Stein and of Perrault and colleagues, the animals were anesthetized, whereas in the other studies, the animals were actively engaged in a fixation task. It was shown (Populin, 2005) that anesthesia affects neuronal sensory responses (modifications of response magnitude, receptive field properties and first-spike latency). Thus, the anesthetized *versus* awake state may explain the differences in bimodal integration between the above studies. Another explanation to sub-additive multisensory integration is the strength of the stimuli: our stimuli were well above perception threshold. Weak stimuli would benefit probably more from multisensory perception, with supra-additivity and enhanced responses (law of inverse effectiveness, Meredith and Stein, 1983).

### Latencies

Our study disclosed median latencies of 89 ms for auditory stimuli, 75 ms for audiovisual stimuli and 141 ms for visual stimuli. In monkeys performing a delayed non-match-to-sample task, Nguyen and collaborators and Van Le and collaborators found latencies between 30 and 500 ms in the pulvinar, which follow a bimodal distribution (Nguyen et al., 2013; Van Le et al., 2013). The mean latencies for the short latency group were 63 ms (Nguyen et al., 2013) and 61 ms (Van Le et al., 2013). The mean latency for the long latency group was 253.5 ms (Van Le et al., 2013). Petersen et al. (1985) measured visual latencies in three parts of the pulvinar (lateral, inferior and dorsomedial). In the dorsomedial pulvinar, they found a mean latency of 84 ms. The latencies in the aforementioned studies are shorter than ours. However, one should notice methodological differences when comparing these latencies. First, our recordings were restricted to the medial pulvinar, whereas in two of the studies mentioned above, recordings were performed either in the whole pulvinar (lateral, medial and inferior (Nguyen et al., 2013)) or in the both the medial and dorsolateral pulvinar (Van Le et al., 2013). Second, our method for latency measurement is different from that used in these studies. Another potential explanation is that, in these studies, the monkeys were performing active tasks in which the visual stimuli were relevant.

Visual latencies were much longer than auditory latencies. This difference has also been observed in the superior colliculus in awake (Populin and Yin, 2002) and anesthetized (Wallace et al., 1996) animals, with comparable or even larger differences (51 ms in the study of Populin and Yin and 83 ms in the study of Wallace and collaborators).

### No threat detection?

One hypothesis about the pulvinar is that it has evolved by selection pressure to allow fast and accurate detection of predators like snakes (Isbell, 2006). This hypothesis is supported by electrophysiological studies showing stronger responses in the pulvinar to snakes and to human or to monkey faces expressing surprise or anger (Maior et al., 2010; Van Le et al., 2013, 2014, 2016). These studies show that the pulvinar detects stimuli providing information about a potential threat, by responding differently and with different latencies depending on the threat level of the stimulus. This hypothesis is also consistent with the potential role of the medial pulvinar in emotion processing (Arend et al., 2015 for a review). Studies have shown alterations of threat recognition in patients with lesions restricted to the medial pulvinar, but not in patients with lesions restricted to the anterior or lateral pulvinar (Ward et al., 2007). Our study cannot confirm this point: in our recordings, neurons responded similarly to snake, macaque and noise stimuli. Sparseness and selectivity indexes revealed a distributed representation, with neurons responding to many stimuli. Latencies also didn’t reveal faster responses for snake stimuli. This discrepancy can be linked to the task: our monkeys were keeping stable fixation while passively viewing or listening to the stimuli. We also repeatedly used only one potentially threatening stimulus, maybe not sufficiently threatening and to which the monkey had probably become accustomed. In the studies mentioned above, the monkeys were involved in an active task, and there were many threatening stimuli among all behaviorally relevant stimuli.

### Functions of the medial pulvinar

The functional role of the pulvinar remains largely unknown, yet some hypotheses have been put forward (Froesel et al., 2021).

First, anatomical studies have shown many sensory connections with the pulvinar (Cappe et al., 2007, 2009a), overlapping territories of projections from different sensory cortices to the medial pulvinar (Cappe et al., 2009a), and projections from the medial pulvinar to the premotor cortex (Cappe et al., 2009b). This suggests that the medial pulvinar is a place of sensorimotor convergence.

Second, studies have shown the involvement of the pulvinar in sensory processing (Almeida et al., 2015; Gattass et al., 1978; Komura et al., 2013; Maior et al., 2010; Mathers and Rapisardi, 1973; Nguyen et al., 2013; Van Le et al., 2013, 2014, 2016; Wilke et al., 2009; Yirmiya and Hocherman, 1987). Some of these studies specified where in the pulvinar single-units were recorded, and demonstrated that the medial pulvinar possesses visual, auditory and somatosensory-responsive neurons. Our study adds to these by showing that the medial pulvinar also contains multisensory audiovisual neurons, with various multisensory integration characteristics.

Third, the pulvinar would be involved in sensory distractor filtering (Fischer and Whitney, 2012) and in visual selection and attention (Saalmann and Kastner, 2009), allowing the detection of important stimuli (behaviorally relevant or requiring fast motor responses for example).

Fourth, it has been hypothesized that the pulvinar plays a major role as mediator or modulator of signal transfer between cortical areas (Benarroch, 2015; Saalmann and Kastner, 2011). Indeed, the replication principle postulates that all directly linked cortical areas would also be indirectly connected *via* the pulvinar (Sherman, 2016, 2017). These cortico-pulvino-cortical pathways would improve the signal-to-noise ratio of information transmitted by modulating the synchrony between these areas (Fries, 2015). These cortico-thalamo-cortical pathways would be faster than cortico-cortical projections and could therefore enable fast multisensory and multisensory-motor interactions between distant cortical regions (Cappe et al., 2009b; Sherman and Guillery, 2002). These two last hypotheses have not been tested specifically for the medial pulvinar.

The medial pulvinar and the amygdala would be part of two emotion processing pathways (the first one, involving also the orbitofrontal cortex, for conscious emotion perception, and the second one, with the superior colliculus, for unconscious emotion perception), both fundamental for survival alore or in complex social groups, by allowing fast behavioral reactions. In addition, the pulvinar may be an important modulator of behavioral flexibility, contributing to the selection of sensory inputs (Froesel et al., 2021). This hypothesis is consistent with the multisensory integration indices in the present study.

All these hypotheses require sensory processing, and ideally multisensory processing. Our study is the first to prove multisensory processing and integration in the pulvinar, more precisely in the medial pulvinar. We propose that the pulvinar combines multiple sources of sensory information to improve the monitoring of the environment, while also playing the role of a general regulation hub for adaptive and flexible cognition.

In future studies, it would be interesting to investigate the same mechanisms with an active task. It would be also of interest to record specifically in the medial pulvinar, with stimuli of different emotion and threat levels, and with stimuli appearing less frequently than the others. This would allow testing the threat detection hypothesis in this structure. If the medial pulvinar is involved in threat detection, the expectation is to have different responses strength and latencies, depending on the stimuli. Another hypothesis that could be tested is the involvement of multisensory integration in sensory distractor filtering and emotion processing. If multisensory integration helps to detect relevant stimuli, there should be enhanced or supra-additive integration for relevant stimuli, but not for irrelevant stimuli.

## Author contributions

C.J. and C.C. designed the study. C.J., C.C. and P.G. executed the experiments. C.J., C.C., P.G. and L.G.N. analyzed the data. A-L. V. C.J., P.G., L.G.N. and C.C. wrote the manuscript. All authors reviewed and edited the manuscript.

## Declaration of interests

The authors declare no competing interests.

## STAR methods

### RESOURCE AVAILABILITY

#### Lead contact

Further information and requests for resources and reagents should be directed to and will be fulfilled by the Lead Contact, C. Cappe.

#### Materials availability

This study did not generate new unique reagents.

#### Data and code availability

The datasets supporting the current study have not been deposited in a public repository because there are still under investigation but are available from the corresponding author on request.

### EXPERIMENTAL MODEL AND SUBJECT DETAILS

#### Animals

Two adult male rhesus monkeys (Macaca mulatta), weighting 7 and 5 kg and 5-years-old, participated to this experiment. They were naive to all procedures. They were housed together at night and separated during the day in cages for food and water delivery. Water was available *ad libitum* (as soon as they were in cage), whereas food (fruits, vegetables and cereals) was controlled and given after each training or recording session. A weight loss criterion of 10% was decided for stopping the experiment, but this never happened. All procedures were approved by the National Committee for Ethical Reflection on Animal Testing in compliance with the guidelines of the European Community on Animal Care (authorization number: 01000.02).

### METHOD DETAILS

#### Experimental procedures

Monkeys were trained to a fixation task. They were sitting in a primate chair (Crist instruments), with the head fixed, in a darkened, sound-attenuated box, in front of a 31 cm-distant screen (BenQ, 60 cm diagonal, 1920 × 1080 pixels, 120 Hz). Two loudspeakers (Creative Gigaworks t20 serie II) were positioned one on each side of the screen. The training took place five days a week, on the morning. The monkeys were food deprived, and the training was their first access to food. Each correct trial was rewarded with diluted compote (0.05 mL/trial).

The task was as follows. To initiate a trial, the monkey had to fixate a 0.5° point (into a square fixation window of 2×2°) for a random time between 500 and 1200 ms. Then a stimulus was presented for 250 ms (auditory, visual or audiovisual stimulus). The monkey had to maintain fixation during stimulus presentation. If fixation was not broken, the monkey was rewarded after a random delay (300-700 ms). Otherwise, the trial was aborted. The intertrial interval was 1000 ms. The task was controlled with EventIDE software (Okazolab Ltd), and eye tracking was performed with an eyetracker (ISCAN ETL 200, Woburn, MA 01801).

A total of nine stimuli was used: three auditory, three visual and three audiovisual stimuli. The auditory stimuli consisted in white noise, in a macaque “coo” call and in a rattlesnake rattle. The auditory stimuli were stereo 44-48 kHz waves normalized to 60 dB, with a 3 ms fading-in and fading-out. Visual stimuli were a picture of random dots, a rhesus macaque head photograph and a rattlesnake picture. All images were normalized in RGB colors with a color depth of 24, sized 453×453 pixels with a 72×72 dpi resolution (final size 19×19°), with a mean luminance of 120 cd/m^2^, and presented in the center of the screen. There was only one exemplar of each stimulus (macaque, snake, noise). The audiovisual stimuli were the synchronous presentation of an auditory and a visual stimulus, always semantically congruent. The stimuli were randomly presented, about 20 times each. A 4×4° white square appeared in the upper right corner of the screen (hidden from the monkey’s view by black tape) at the same time as the stimulus presentation, to precisely indicate onset and offset of the stimulus.

#### Surgical procedures

A first surgery was performed to implant an MRI-compatible headpost (Crist Instrument) to the skull. After one month of recovery, the monkeys were trained to the fixation task. When they reached 90-95% correct trials, a second surgery was performed to stereotaxically implant a footed stainless-steel recording chamber (Crist Instrument) above the right somatosensory cortex (S2) (centered on average AP = 6.5 and ML = 0.75). The skull within the chamber was removed to allow direct access to the brain. The dura mater was left intact and was protected by a sterile silicone patch.

All surgeries were performed under anesthesia, induced with a mixture of tiletamine/zolazepam (Zoletil 50®, 5 mg/kg) and glycopyrrolate bromide (Robinul®, 0.01 mg/kg), and maintained with isoflurane (1.5%) after intubation. Analgesics were administered (Tolfedine 4 mg/kg and buprenorphine chlorhydrate (Vetergesic®) 0.01 mg/kg) during surgery and the following days. An antibiotic treatment (Amoxicillin (Clamoxyl^®^ LA) 15 mg/kg) was administered during the first week.

The location of the recording chamber was determined before surgery by comparing stereotaxic 3T anatomical MRI scans to sections of the stereotaxic atlas of the brain of Macaca mulatta (Paxinos et al., 2000; Saleem and Logothetis, 2007). After implantation, the locations of the recording sites were confirmed by comparing the atlas sections to stereotaxic anatomical MRI scans of the monkeys’ head with a tungsten microelectrode inserted at target coordinates. The locations of the recording sites are relative to the zero coordinates defined in the stereotaxic atlas.

#### Recordings

Neuronal activity was recorded with tungsten microelectrodes (5-7 MΩ at 1 kHz, Frederick Haer Company, Bowdoinham, ME). Electrodes were inserted daily into the medial pulvinar (ML between 4.9 and 9.8 mm, AP between 5 and 9.8 mm, depth between 17.9 and 23.8 mm below the cortical surface). Electrode insertion was performed with an oil hydraulic micromanipulator (Narishige MO-972) attached to the recording chamber.

Recorded signal was sampled at a frequency of 40 kHz with a 1401 power acquisition interface (CED, Cambridge, UK), amplified with a gain of 1000 (NL104) and band-pass filtered between 200 Hz and 8 kHz (NL125) by a Neurolog system (Digitimer, Hertfordshire, UK). The 50 Hz noise was eliminated with a Humbug device (Digitimer). Signal was displayed and recorded with Spike2 software (CED). Data were analyzed with scripts in Spike2 software environment.

#### Unit activity

Collected data usually contained spikes from multiple units. Spike sorting was performed with the PCA-based cluster analysis of Spike2 software. For each cluster, the refractory period was determined with an inter-spike interval histogram. Spikes were considered issued from a unique neuron (single-unit) if the refractory period was ≥ 1.2 ms.

### QUANTIFICATION AND STATISTICAL ANALYSIS

#### Neurons’ classification

Only neurons with a large enough activity (more than 800 spikes on the whole recording period) were analyzed. To determine if there was a response to the stimuli, we first calculated, for each stimulus presentation, the mean firing rate during stimulus presentation (0-250 ms) and during the baseline period (−500-0 ms). The distributions of the ≈20 values of spontaneous and evoked firing rates obtained for each type of stimulus were then compared with the Wilcoxon signed-rank test. A p-value of 0.05 was used as a threshold to indicate the presence of significant excitatory or inhibitory responses. Responses to different stimuli were compared using the Kruskal-Wallis test and the Mann-Whitney test applied to the distributions of firing rates calculated for each stimulus presentation. Again, the threshold p-value was 0.05, and a Bonferroni correction was applied for multiple comparisons. This allowed to classify neurons in four categories: visual (response to visual stimuli only), auditory (response to auditory stimuli only), classic multisensory (response to audiovisual stimuli only, or response to both auditory and visual stimuli, or response to both unisensory and multisensory stimuli with different response strengths to these stimuli, or response only to unisensory stimuli with response different from that to the audiovisual stimuli) and complex multisensory neurons (response to auditory or visual or audiovisual stimuli, depending on the category of the stimuli).

#### Neurons’ responses characterization

As indicated in the results for more readibility, we used four indexes to characterize the responses of the neurons. Two indexes were calculated to characterize the multisensory integration of multisensory neurons: the additivity index and the amplification index. When neurons were multisensory for more than one stimulus category, indexes were averaged.

Two other indexes were computed for all neurons: the selectivity index and the sparseness index. The selectivity index shows whether a neuron was selective for one stimulus (index close to 1) or equally responsive to different stimuli (index close to 0). The sparseness index indicates whether the neuron responded similarly to all stimuli (index close to 1) or whether it responded exclusively to one or few stimuli (index << 1).

#### Latencies

Latencies were calculated using the half-rise latency method (Levick, 1973; Nowak et al., 2010). To improve the temporal resolution while preserving a good signal-to-noise ratio, 10 peri-stimulus time histograms with a bin width of 10 ms were computed with a 1 ms offset increment for each calculation. The 10 PSTHs were then averaged with a 1 ms resolution. The peak amplitude and the half-peak amplitude were determined. Only peaks with a half-peak amplitude higher than the mean + 3 SD in PSTH prestimulus baseline (−500 – 0 msec) were considered. The time after stimulus onset at which the half-peak amplitude is first reached corresponds to the half-rise latency. A further constraint was that the amplitude remained ≥ half-peak height for at least 7 consecutive ms. Note that this method, which requires assessment of significance over the time course of the response, is more constrained than that used for determining presence of significant response over the whole stimulus presentation (“neurons classification” above). This explains why the number of neurons with significant latencies is less than the number of neurons with significant responses.

## Notes

### Competing Interest Statement

The authors have declared no competing interest.

