## Supplementary figures and images for "Multisensory integration in neurons of the medial pulvinar of macaque monkey"

### Supplemental Figure 1

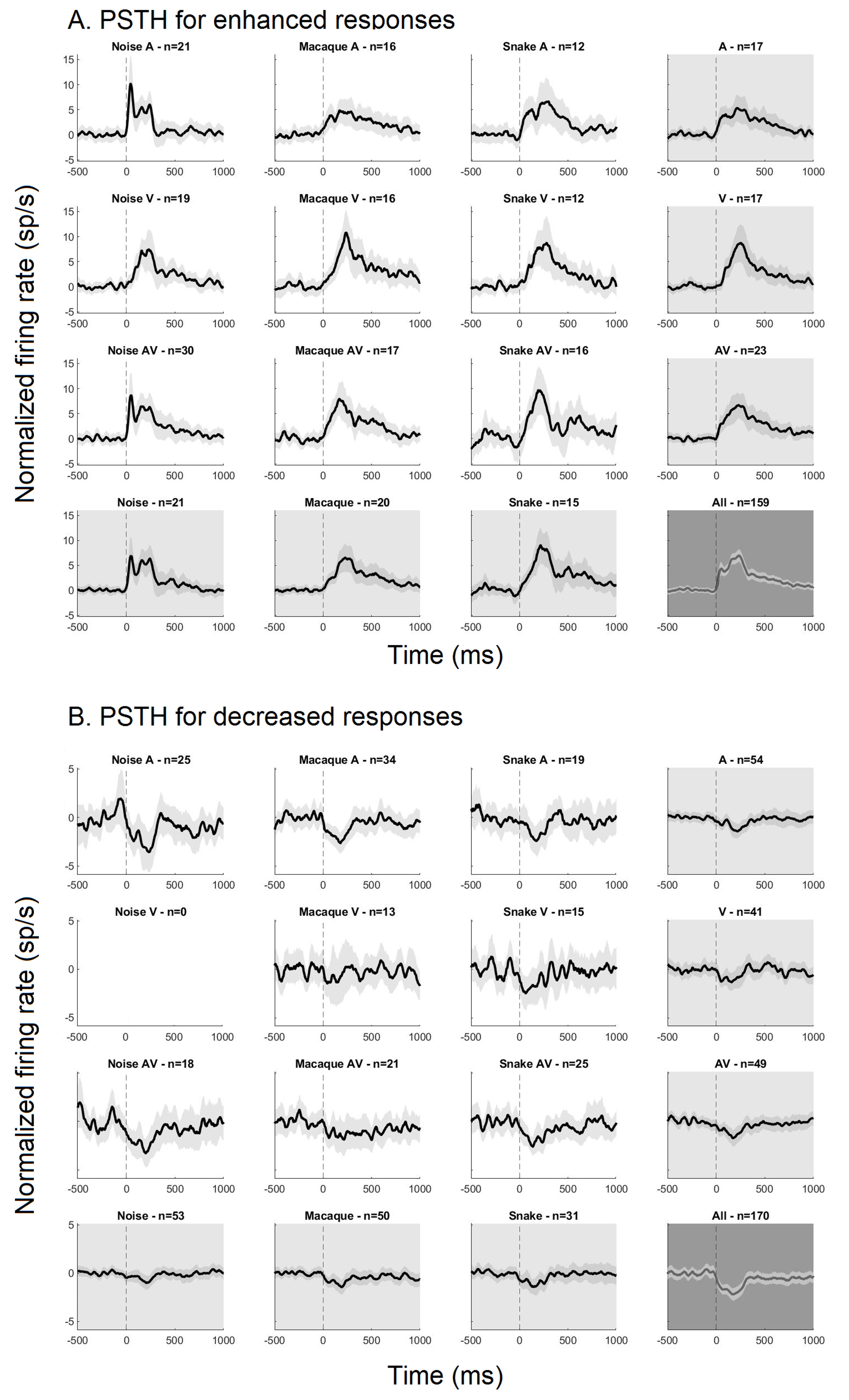
