## Supplemental Figure 1 - Legend for "Multisensory integration in neurons of the medial pulvinar of macaque monkey"

**Figure S1: Global responses of neuronal populations according to the response types**

Global PSTHs represent the mean firing rate (sp/s) of neuronal populations according to the responses types, for each stimulus and modality: enhanced (A) or decreased (B) responses. PSTHs rows correspond, from top to bottom, to auditory, visual, audiovisual and pooled modality conditions. PSTHs columns correspond, from left to right, to noise, macaque, snake and pooled stimuli conditions.

Responses were averaged over all neurons showing a statistically significant enhancement (A) or suppression (B) for each stimulus, without (white background) or after (grey background) pooling across stimuli ($4^{th}$ row) or modalities ($4^{th}$ column).

The PSTHs with the dark grey background represent population average for all enhanced or all decreased responses (same as fig. 2A, B).

Black line corresponds to the mean and gray shading to ± 95% CI.
